# GTFtools: a Python package for analyzing various modes of gene models

**DOI:** 10.1101/263517

**Authors:** Hong-Dong Li

## Abstract

**Summary:** Gene-centric bioinformatics studies frequently involve calculation or extraction of various features of genes such as gene ID mapping, GC content calculation and different types of gene lengths, through manipulation of gene models that are often annotated in GTF format and available from ENSEMBL or GENCODE database. Such computation is essential for subsequent analysis such as intron retention detection where independent introns may need to be identified, converting RNA-seq read counts to FPKM where gene length is required, and obtaining flanking regions around transcription start sites. However, to our knowledge, a software package that is dedicated to analyzing various modes of gene models directly from GTF file is not publicly available. In this work, GTFtools (implemented in Python and not dependent on any non-python third-party software), a stand-alone command-line software that provides a set of functions to analyze various modes of gene models, is provided for facilitating routine bioinformatics studies where information about gene models needs to be calculated.

**Availability:** GTFtools is freely available at www.genemine.org/gtftools.php

## 1. Introduction

Computing or extracting various features of genes such as gene ID mapping, identification of independent introns that do not overlap with any exons of any transcripts^1,2^, and calculation of different types of gene lengths, is frequently confronted in routine bioinformatics analysis. Such computation is essential for subsequent analysis such as alternative splicing analysis including intron retention detection where independent introns need to be identified^1,2^, obtaining flanking regions around transcription start sites (TSS)^3^, and gene expression normalization by converting RNA-seq read counts to FPKM where exonic gene length needs to be calculated^4^. A popular tool that can accomplish part of these tasks is BioMart (https://www.ensembl.org/biomart). However, it is dependent on database querying and can be slow sometimes. Also, users need to be familiar with the field names of tables in the back-end database of BioMart, which may be inconvenient.

Since gene model annotation is the basis of analyzing various modes of gene features, an efficient way for such analysis would be directly interrogating gene models, which are often annotated in GTF files and available from ENSEMBL or GENCODE^5^. Here, GTFtools, a stand-alone command-line software that provides a set of functions to extract features from gene models, is presented. It is not dependent on any existing bioinformatics tools and would thus be easy to install and use. GTFtools provides a new and important tool for facilitating routine bioinformatics analysis.

## 2. Implementation

GTFtools is implemented in Python. Command-line options are parsed by the ‘argparse’ package, which hence needs to be installed. GTFtools takes a GTF file (ENSEMBL or GENCODE) as input, and output what is specified by users in bed or bed-like format. Correctness of bed file operation such as merging and subtracting was validated by Bedtools^6^. The major functions currently implemented include the calculation of merged exons, independent exons^2^, four types of gene length (the mean, median and max of lengths of isoforms of a gene, and the length of merged exons of isoforms of a gene), UTR, TSS, gene symbol-ID mapping etc. Details are shown in **Table 1**. Coupled with other software like Bedtools^6^, sequence features such as GC content and dinucleotide frequency can be further analyzed.

**Table 1.** Major functions in the GTFtools (version 0.6.0).

| Options | Functions | Notes |
| --- | --- | --- |
| -m | For each gene, calculate **non-overlapping exons** by merging exons of all splice isoforms from the same gene. The output is merged exons in bed format. | Merge all exons of all isoforms of the same gene. |
| -d | Calculate **independent introns** which is defined as introns (or part of introns) that do not overlap with any exons of any genes in the genome<sup>2</sup>. It is calculated by subtracting merged exons from genes. The output is in bed format. | Useful for intron retention detection. |
| -l | Calculate gene lengths. Since a gene may have multiple isoforms, gene length calculation is not as simple as it appears. The mean, median and maximum of the lengths of isoforms for a given gene are considered. In addition, the length of merged exons of all isoforms (i.e. non-overlapping exon length) is also calculated. So, in total, **four different types of gene lengths** (the mean, median and max of lengths of isoforms of a gene, and the length of merged exons of isoforms of a gene) are provided. | Needed for e.g. calculating FPKM in RNA-seq data analysis, where gene length is required. |
| -g | Output gene coordination and ID mappings in bed format. |  |
| -s | Output isoform coordination and parent-gene IDs in bed format. |  |
| -e | Output exons in bed format. |  |
| -i | Output introns in bed format. |  |
| -k | Calculate introns (part of introns) that overlap with exons of other isoforms. The output is in bed format. |  |
| -u | Output UTR in bed format. |  |
| -t | Out transcription start site (TSS)-flanking region in bed format. |  |
| -c | Specify chromosomes to analyze. Dash(-) and comma(,) are allowed. For example, '-c 1-5,X,Y' indicates 7 chromosomes: 1 to 5 together with X and Y. Default is 1-22, X and Y. |  |

## 3. Conclusion

A python package for computing or extracting various modes of gene models in GTF format has been developed in this work. It takes GTF files downloaded from the ENSEMBL or GENCODE database as input, and outputs what is specified by users in bed or bed-like format. It is expected that GTFtools would become a useful tool and facilitate routine bioinformatics analysis.

